# A streamlined method to obtain biologically active TcdA and TcdB toxins from *Clostridioides difficile*

**DOI:** 10.1101/2023.07.19.549664

**Authors:** Afi Akofa Diane Sapa, Anaïs Brosse, Héloïse Coullon, Gauthier Pean de Ponfilly, Thomas Candela, Alban Le Monnier

**Affiliations:** Université Paris-Saclay, INRAE, AgroParisTech, Micalis Institute, 78350, Jouy-en-Josas, France; Service de Microbiologie clinique, GH Paris Saint-Joseph, Paris, France

**Keywords:** *Clostridioides difficile*, toxin, recombinant protein, cytotoxic activity, TcdA, TcdB

## Abstract

The major virulence factors of *Clostridioides difficile* (*C. difficile*) are enterotoxin A (TcdA) and cytotoxin B (TcdB). The study of toxins is a crucial step in exploring the virulence of this pathogen. Currently, the toxin purification process is either laborious and time-consuming in *C. difficile* or performed in heterologous hosts. Therefore, we propose a streamlined method to obtain functional toxins in *C. difficile*. Two *C. difficile* strains were generated each harboring a sequence encoding a His-tag at the 3’ end of *C. difficile* 630Δ*erm tcdA* or *tcdB* genes. Each toxin gene is expressed using the P_tet_ promoter inducible by anhydro-tetracycline. The purification yields were estimated to be 0.28 mg per liter and 0.1 mg per liter for rTcdA and rTcdB, respectively. In this study, we successfully developed a simple routine method that allows the production and purification of biologically rTcdA and rTcdB active toxins with similar activities compared to native toxins.

## Introduction

*Clostridium difficile* (*C. difficile*) is a Gram-positive, anaerobic, spore-forming, toxin-producing bacillus that was renamed *Clostridioides difficile* in 2016 [1]. *C. difficile* was initially identified as part of the flora of healthy infants in 1935 [2]. It was subsequently described as a causative agent of digestive tract infections often involved in antibiotic-associated diarrhea ranging from mild to severe and life-threatening complications such as pseudomembranous colitis. *C. difficile* infections (CDI) are associated with high recurrence rates, reaching up to 30% of cases [3–5]. CDI have been recognized as a leading cause of healthcare-associated infections and more generally, a substantial threat to public health [6–8]. The main risk factor for developing CDI is antibiotic therapy [9] which can disrupt the gut microbiota. In this context of dysbiosis, spores of *C. difficile* will germinate into vegetative forms which will then colonize the gut microbiota, multiply, and produce virulence factors, in particular enterotoxin A (TcdA) and cytotoxin B (TcdB) that lead to the development of symptoms. TcdA and TcdB are encoded by the *tcdA* and *tcdB* genes, localized in the 19.6kb pathogenesis locus (PaLoc), along with three additional genes that allow regulation and release of the toxins (*tcdR*, *tcdC,* and *tcdE,* [10]). TcdA and TcdB are expressed when bacterial cells enter the stationary phase, likely caused by the limitation of nutrients [11]. TcdA and TcdB belong to the family of Large Clostridia Toxin (LCTs) due to their high molecular weight (308 and 270 kDa respectively). The LCTs can induce profound changes in cell morphology [12]. These toxins are composed of four functional domains; the N-terminal glucosyl-transferase region, the auto-processing cysteine protease domain, a membrane translocation region (CROPS), and a C-terminal Receptor-Binding Domain region. After binding to their receptors on the cell surface, endocytosis is initiated and toxins are internalized into endosomes. Subsequent acidification of the endosomes induces a conformational change of the delivery domain, resulting in pore formation and translocation of a catalytic glucosyltransferase domain across the endosomal membrane. The toxins subsequently undergo autocleavage which releases the N-terminal glucosyltransferase region into the cytoplasm [13]. The glucosyltransferase activity domain targets and inactivates Rho GTPases, including Rac1 and Cdc42 at Thr35 as well as RhoA at Thr37. Inactivation of Rho GTPases leads to actin-depolymerization, which results in cell rounding, disaggregation of the actin cytoskeleton, loss of intestinal epithelial barrier function and cell death. In addition to their direct cytotoxic effects, TcdA and TcdB elicit a pro-inflammatory response contributing to tissue damage and leading to severe complications such as pseudomembranous colitis [14,15]. In recent years, studies have provided further insight into the structure of TcdA and TcdB across *C. difficile* strains, leading to the identification of distinct subtypes of toxins [16]. Of note, clade 2 hypervirulent *C. difficile* strains have been linked to the expression of toxin subtypes TcdB2 and TcdB4. Importantly, subtypes of TcdB toxins have also been linked to the recognition of distinct receptors within the host [17,18]. Taken together, these recent results highlight the need for adequate tools for accurate toxin studies.

Having biologically functional purified toxins is an important tool to understand these toxins’ activities in the modulation of the pathogenesis of CDI, the host immune response and more importantly for the development of CDI treatments. In addition, clinical research relies on *C. difficile* toxins to quantify anti-TcdA or TcdB antibodies by ELISA or for neutralizing antibody tests. The native toxins are usually purified from toxigenic *C. difficile* VPI 10463 culture supernatant [19–21] but the purification process is fastidious, time-consuming, and requires multiple steps. In this study, we aimed to develop a simplified method for the purification process of biologically active native *C. difficile* TcdA and TcdB by directly using *C. difficile* toxigenic strain 630Δ*erm* as the host.

## Materials and methods

### Bacterial strains, plasmids, and growth conditions

Bacterial strains, primers, and plasmids used in this study are detailed in S1, S2 and S3 Tables. *E. coli* strains were grown aerobically at 37°C in Luria Bertani (LB) medium (Becton, Dickinson) supplemented with Ampicillin (Amp; 100 µg.mL^-1^) or Chloramphenicol (Cm; 25 µg.mL^-1^) as appropriate. *C. difficile* strains were grown at 37°C in an anaerobic chamber using Brain Heart Infusion (BHI) medium (Becton, Dickinson) or tryptone-yeast medium (TY; 3 % Bacto tryptose (Becton, Dickinson), 2% Bacto yeast extract (Becton, Dickinson), and 0.1 % thioglycolate, adjusted to pH 7.4) supplemented as appropriate with Thiamphenicol (Tm; 15 μg.mL^-1^), Aztreonam (Az; 16 μg.mL^-1^) or nonantibiotic analog Anhydro - tetracycline (ATc; 0.05 µg.mL^-1^, 0.1 µg.mL^-1^, 0.25 µg.mL^-1^, 0.5 µg.mL^-1^).

### Constructs and cloning in *C. difficile* 630Δ*erm* strain

Plasmid extraction (Omega, VWR), endonuclease digestion (New England Biolabs), ligation (New England Biolabs), and agarose gel electrophoresis were carried out as described by Maniatis *et al* [22]. The Phusion polymerase (New England Biolabs) used for Polymerase Chain Reaction (PCR) and Golden gate cloning kit (New England Biolabs) were carried out according to the manufacturer’s instructions. All inserts cloned in the constructed plasmids were sequenced (Eurofins Genomics).

In order to have a plasmid suitable for the Golden gate cloning strategy, the pTC130 was constructed. First, the spectinomycin cassette from the pAT28 plasmid [23] was amplified using primers 2836/2837 and cloned into the pBlunt2 plasmid (Invitrogen), giving rise to pTC129. The 1.09 kb BamHI/EcoRV fragment extracted from pTC129 was ligated into pMSR [24] digested with BamHI/PvuII, giving rise to pTC130.

Plasmids pADS1 and pADS2 were constructed for the genomic insertion of 6xHis-tags before the STOP codons of *tcdA* and *tcdB* respectively, using the following strategy. Two DNA fragments flanking the STOP codon of *tcdA* were obtained by PCR using *C. difficile* 630Δ*erm* genomic DNA and primers 2854/2851 and 2852/2853. These DNA fragments were cloned into pTC130 by the golden gate method [25] giving rise to pADS1. Similarly, two PCR DNA fragments flanking the STOP codon of *tcdB* were obtained using 2858/2859 and 2860/2861 primer pairs on *C. difficile* 630Δ*erm* genomic DNA and cloned into pTC130 by the golden gate method, giving rise to pADS2. Plasmids pADS1 and pADS2 were confirmed by enzymatic restriction as well as sequencing of the inserts. pADS1 and pADS2 recombinant pseudo suicide plasmids were transferred by heterogramic conjugation from conjugative *E. coli* strain HB101 (pRK24) to *C. difficile* 630Δ*erm*. The recombinant strains AD1 and AD2 were selected as described by Peltier *et al*. [24]. Recombinant mutants were confirmed by PCR and sequencing of the targeted genomic region.

The same strategy was used to build the pADS3 and pADS4 plasmids to replace the *tcdA* and *tcdB* promoters with a P_tet_ promoter. Homologous fragments upstream (CO1) and downstream (CO2) of the *tcdA* promoter were amplified from *C. difficile* 630Δ*erm* extracted genomic DNA using primers 3046/3047 and 3050/3051. The P_tet_ promoter from the pRPF185 plasmid [26] was amplified using primers 3048/3049. These three fragments were cloned into pJV7 [27] by the golden gate method, giving rise to pADS3. Similarly, PCR DNA fragments CO1 and CO2 flanking the promoter of *tcdB,* obtained using primers 3052/3053 and 3056/3057 on *C. difficile* 630Δ*erm* genomic DNA, as well as the P_tet_ promoter, amplified from the pRPF185 plasmid using primers 3054/3055, were cloned into pJV7 by the golden gate method, giving rise to pADS4. After confirmation by enzymatic restriction and sequencing, the pADS3 and pADS4 recombinant pseudo-suicide plasmids were transferred by heterogramic conjugation from conjugative *E. coli* strain HB101 (pRK24) to *C. difficile* strains AD1 (*tcdA-*His) and AD2 (*tcdB-*His). Selection of the recombinant strains AD3 (P_tet_*-tcdA-*His) and AD4 (P_tet_*-tcdB-*His) was performed as described by Peltier *et al*. [24]. Recombinant strains were subjected to PCR screening and sequencing to confirm the correct insertion of the fragments and the absence of additional mutation. Strains and plasmids detailed in this study are available for researchers upon request.

### Expression and purification of recombinant toxins

#### From AD1 and AD2 strains

Cultures of *C. difficile* strains AD1 and AD2 were prepared in 600 mL of TY medium, inoculated at a 600 nm optical density (OD_600nm_) of 0.05 using overnight cultures, and allowed to grow for 75 to 92 h in an anaerobic chamber at 37°C to induce toxin release in nutrient limiting medium [11]. Bacteria were centrifuged and culture supernatants were precipitated with 45 g of ammonium sulfate for 100 mL of bacterial culture followed by shaking at room temperature for 5 min. Precipitates were centrifugated (12200 g) for 10 min at 4 °C, and the pellet was resuspended in 40 mL of denaturing buffer (8 M Urea, 0.1 M NaH2PO4, 0.01 M Tris-HCl, pH 8). pH was adjusted to 8 after resuspension to allow the binding of 6x His-tagged toxins to Ni-NTA agarose resin (reference: 30210; Qiagen). Resulting samples were passed through chromatography columns containing the Ni-NTA agarose resin and were eluted with denaturing buffer containing increasing concentrations of Imidazole (10 mM, 20 mM, 30 mM, 40 mM, 60 mM, 80 mM, 100 mM, 200 mM, and 1 M). For analysis, fractions were loaded into a sodium dodecyl-sulfate polyacrylamide gel electrophoresis (SDS-PAGE) gel using 6% stacking gel and 10% separation gel followed by a standard Coomassie staining.

#### From AD3 and AD4 strains

Cultures of *C. difficile* strains AD3 and AD4 were prepared in 1000 mL, and 2000 mL of BHI medium respectively, inoculated at OD_600nm_ of 0.05 using overnight cultures, and incubated in an anaerobic chamber at 37°C. Cultures were allowed to grow until reaching an OD_600nm_ of 0.5 before toxin expression was induced by the addition of ATc (0.5 µg.mL^-1^ for recombinant toxin A rTcdA or 0.1 µg.mL^-1^ for recombinant toxin B rTcdB). Cultures were incubated for 4 h after ATc addition, bacteria were centrifuged and pellets were frozen at −20°C. Cell pellets were resuspended in 40 mL of lysis buffer (50 mM NaH_2_PO_4_, 300 mM NaCl, 10 mM Imidazole, pH 8). For lysis, bacteria were then incubated for one-hour at 37 °C, followed by a brief sonication to reduce viscosity. Lysates were centrifuged (12200g) for 30 min at 4°C. Lysate supernatants were then passed through chromatography columns containing Ni-NTA agarose resin (reference: 30210; Qiagen). The bound 6x His-tagged toxins were eluted with elution buffer (50 mM NaH2PO4, 300 mM NaCl, 200 mM Imidazole, pH 8). The eluents were desalted using a PD10 column (reference 17085101; Cytiva), and the recombinant toxins were eluted with 50 mM NaH_2_PO_4_, and 300 mM NaCl, and conserved at −20°C. Samples were analyzed by SDS-PAGE as previously described. Purified proteins were obtained at 80 µg.mL^-1^ and 60 µg.mL^- 1^ for rTcdA and rTcdB respectively on average.

### Cytotoxicity assay

Cytotoxic activity of purified recombinant toxins rTcdA and rTcdB was determined using Vero (African green monkey kidney) cell monolayer culture with a protocol adapted from Cartman *et al* [28]. Briefly, cells were seeded in a 96-well plate (TPP) with 100 µL cell suspension at the density of 2×10^5^ cells mL^-1^ in Dulbecco’s modified Eagle’s medium (DMEM) (GIBCO) supplemented with 10% (v/v) fetal calf serum (Dutscher). The plates were incubated for 24 h at 37°C, 5 % CO_2_ to obtain the cell monolayer before adding the toxins. In order to evaluate the cytotoxicity of the recombinant toxins compared to native TcdA and TcdB, cells were treated with either TcdA, TcdB, rTcdA or rTcdB using four different concentrations ranging from 4 µg.mL^-1^ to 32 ng.mL^-1^. These dilutions were obtained by preparing fivefold serial dilutions of toxin stocks, in DMEM medium supplemented with 0.1 % of Bovine serum albumin solution (BSA) (Sigma). In addition, cells in DMEM and 0.1% of BSA only were included as a negative control. The *C. difficile* native toxins TcdA and TcdB used in this study were produced in *C. difficile* VPI 10463 strain and generously gifted by Pr Michel Popoff (Pasteur Institute, Paris) [19,21]. After 18 h of incubation, morphological alterations of Vero cells were observed under phase contrast microscopy (Zeiss). Results presented here are representative of at least two independent experiments.

### Disruption of the actin cytoskeleton by purified recombinant toxins

A monolayer of Vero cells was obtained by seeding 1 mL of a 1×10^5^ cells mL^-1^ suspension per well in a 24-well plate (TPP), followed by incubation for 24 h at 37°C, 5 % CO_2_. Sub-confluent cells were then incubated for 18h with 4 µg.mL^-1^ of native toxins (positive controls) or purified recombinant toxins. In addition, a negative control was also added by keeping cells in DMEM and 0.1% of BSA only. After incubation, cells were fixed and stained with Rhodamine Phalloidin (reference: P1951, Sigma) and DAPI contained in the Fluoromount-G^TM^ mounting medium (reference: 00495952, Invitrogen). Five random fields were acquired for each condition, and FIJI software was used to determine the mean area per cell [29]. For each field randomly selected, using a thresholding method and particle analysis, we used the actin channel to determine the area of the field occupied by cells, expressed in µm^2^ and the DAPI channel to count the number of nuclei in the field. We then expressed the average cell area, expressed in µm^2^/cell, by dividing the total cell area by the number of cells. Finally, additional images were acquired with the x63 objective in order to illustrate cell morphological alterations induced by toxins.

### Comparison of native or recombinant toxins for the quantification of serum antibodies by quantitative ELISA

The TcdA and TcdB antibody titers were determined by quantitative enzyme-linked immunosorbent assay (ELISA) using a protocol adapted from Péchiné *et al*. [30]. In order to allow quantitative measurements of serum antibody titers, we added a calibration range by serially diluting a polyclonal IgG antibody from human sera solution (reference: I2511, Sigma), to reach concentrations ranging from 0.68 ng.mL^-1^ to 22 ng.mL^-1^. Briefly, ELISA microplates (MaxiSorp Nunc, reference: 439454) were coated with 1 µg.mL^-1^ of native or purified recombinant toxins in a coating buffer (PBS – Deoxycholate 0.1%) and incubated overnight at 35°C. Microplates were washed four times by adding 250 µL of PBS-Tween 0.1% in each well. After this step, wells were incubated with 200 µL of a blocking buffer (PBS - Bovine Serum Albumin 3% (BSA reference: A9418, Sigma)) during one hour at 35°C. After washing the blocking buffer, serum samples from *C. difficile* recovered patients were tested in technical duplicates at serial dilutions from 1/20 to 1/2560 in dilution buffer (PBS – Tween 0.1% - BSA 0.1%). After a washing step, plates were incubated with a secondary antibody (anti-Human IgG-Phosphatase alkaline antibody (reference: A9544, Sigma)), incubated for one hour at 35°C, and washed four times with PBS-Tween 0.1%. Finally, revelation was done by adding the p-nitrophenyl phosphate substrate (reference: N2770, Sigma) to each well. The light-absorption was measured at 450 nm and anti-TcdA or anti-TcdB IgG concentrations were determined using calibration ranges. Serum samples used in these assays were obtained from seven CDI patients from the SERODIFF study (ClinicalTrials.gov Identifier: NCT01946750). For the SERODIFF study, participants were recruited from December 1^st^ 2012 to June 30^th^ 2017. Written informed consent forms were collected for each participant.

### Neutralization antibody assay

For the neutralization antibody assay, 96 well plates (TPP) were prepared to obtain monolayers of Vero cells using the procedure described in the cytotoxicity assay section. Antibodies used in these assays were obtained from serum samples collected from a CDI patient included in the SERODIFF study. In order to confirm that the recombinant toxins were also susceptible to the neutralizing activity of these antibodies, 75µL of a 4 µg.mL^-1^ solution of rTcdA or rTcdB was mixed with 75 µl of 1/10 dilution of serum from a CDI patient and incubated at 37 °C for 60 min [31,32]. The mixtures were then added to monolayers of Vero cells and the plates were incubated in 5 % CO_2_ at 37 °C for 18 h. A negative control (cells in DMEM and 0.1% of BSA only) and positive controls (4 µg.mL^-1^ and 0.8 µg.mL^-1^ of rTcdA or rTcdB alone) were also added. Cell morphological alterations were observed under phase contrast microscopy (Zeiss).

### Phylogenetic analysis

Phylogenetic analyses were conducted on the following 39 genomes of *C. difficile* strains: 630Δ*erm*, A685, CD015, CD196, CIP1079324, E1, E7, E12, E14, VPI 10463, E15, E16, E23, E24, E25, E28, NAP07, NAP08, QCD_23m63, QCD_32g58, QCD_37x79, QCD-63q42, QCD_66c26, QCD_76w55, QCD_97b34, T3, T5, T6, T10, T11, T15, T17, T19, T20, T22, T23, T42, T61 and R20291. Selected genomes and protein sequences were obtained from Marc Monot [33]. In addition, protein sequences of large clostridial toxins from *Paeniclostridium_sordellii* were obtained from NCBI, for TcsH (AGK40891.1) and TcsL (Q46342.1). See supplementary Table S4 for genome accession numbers.

For phylogenetic trees, protein sequences were first aligned with MUSCLE provided by EMBL-EBI (http://www.ebi.ac.uk/Tools/msa/muscle/) [34]. Phylogeny reconstruction was performed with FastTree (version 2.1.11) using a local installation and running with default settings [35]. Generated trees were exported and uploaded to iTOL (version 6, https://itol.embl.de/) for visualization [36] and bootstrap metadata was visualized using colored tree branches (red = 0 / blue = 1). Sequence alignments obtained in MUSCLE were visualized using BioEdit [37] to generate the amino acid alignment comparisons for TcdA and TcdB in *C. difficile* strains 630Δ*erm*, R20291 and VPI 10463.

### Statistical analysis and software

Analyses were performed using GraphPad Prism 9 (GraphPad Software, San Diego, California, USA). Linear correlation was determined by Pearson correlation coefficient. For microscopy, figure assembly was done using the FigureJ plugin [38].

## Results

### Selection of *C. difficile* strain

Since previous studies relied on the VPI 10463 strain for the purification of TcdA and TcdB, we first sought to determine whether the native toxins from the 630Δ*erm* strain were structurally comparable to the toxins from the VPI 10463 strain. We performed a protein sequence comparison for TcdA and TcdB across 39 *C. difficile* strains and generated phylogenetic trees of each protein list (Fig 1, panels A and B). Each protein entry was subsequently annotated based on their described toxin subtypes [16]. Analysis of these trees showed a distribution of TcdA proteins in tree distinct clusters, which appeared consistent with the toxin subtypes (Fig 1A; TcdA1, TcdA2, TcdA3). Similarly, TcdB proteins appeared in three distinct clusters, consistent with three TcdB subtypes (Fig 1B; TcdB1, TcdB2, TcdB5). In these trees, the TcdA and TcdB proteins of the 630Δ*erm* and VPI 10463 strains were found in the same clusters, suggesting that they are of the same toxin subtype (TcdA1 and TcdB1 respectively). In addition, percent identity matrixes showed identity scores of 99.82% and 100% for TcdA and TcdB respectively (S1 File). Of note, the full-length alignment of TcdA_630Δ*erm*_ and TcdA_VPI 10463_ showed only two differing residues between both proteins at position 2421-2422, which corresponds to the CROPS domain associated with membrane translocation (Fig 1C and S1 File). Taken together, these results demonstrate that the toxins of the 630Δ*erm* are nearly identical to those of the VPI 10463 strain, confirming the potential use of the 630Δ*erm* as a replacement of the VPI 10463 strain for toxin production.

**Fig 1.**
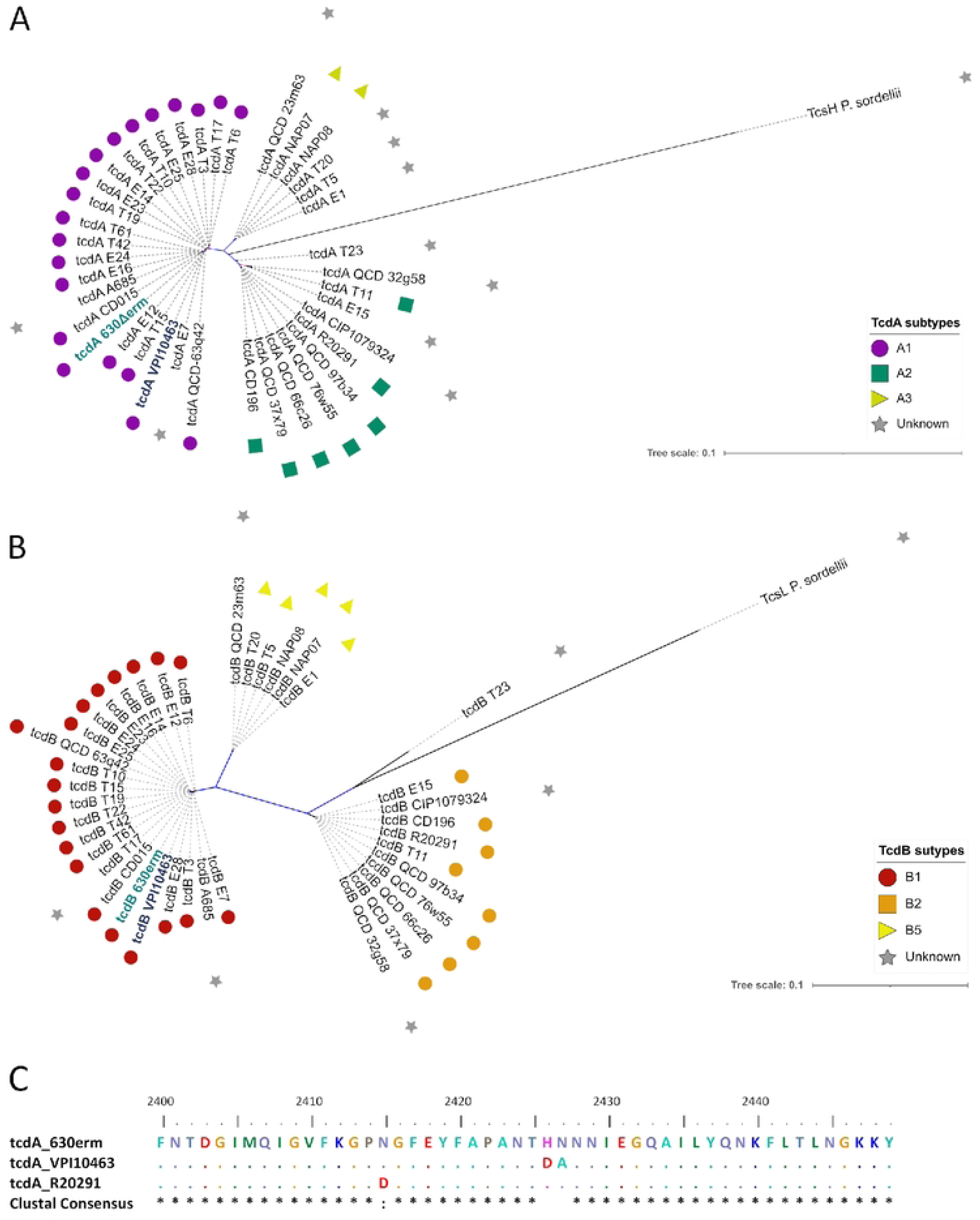
Toxins of the 630Δ*erm* and VPI strains are found in shared clusters. Unrooted phylogenetic trees of (A) TcdA and (B) TcdB proteins obtained from 39 *C. difficile* genomes, with bootstrap values represented as branch colors ranging from 0 in red to 1 in blue. Symbols represent the toxin subgroups for each protein in the trees: TcdA1 = purple circle / TcdA2 = green square / TcdA3 = yellow triangle / TcdB1 = red circle / TcdB2 = orange square / TcdB5 = yellow triangle. In both graphs, gray stars represent proteins for which the toxin subtype is either unknown (*C. difficile* proteins) or not applicable (TcsH and TcsL). Reference sequences from the *C. difficile* 630Δ*erm* and VPI 10463 strains are indicated in bold letters. In addition, each tree contains the closest Large Clostridial Toxin for the respective toxin (TcsH from *P. sordellii* for TcdA / TcsL from *P. sordellii* for TcdB). (C) Multiple sequence alignment of positions 2400 to 2450 of TcdA proteins using *C. difficile* 630Δ*erm*, VPI 10463 and R20291 strains, with the resulting clustal consensus sequence. Identical amino acids are represented as dots.

### Expression and purification of recombinant toxins

We then constructed two pseudo-suicide plasmids pADS1 and pADS2. These plasmids were used to transfer a sequence encoding a His-tag at the 3’ end of *tcdA* and *tcdB* genes by conjugation and generate *C. difficile* strains AD1 and AD2 (S1 Fig), respectively for the production of recombinant toxins rTcdA and rTcdB. The resulting AD1 (*tcdA-*His) and AD2 (*tcdB-*His) strains were grown in TY medium during 75 to 92 h to induce *tcdA* and *tcdB* expression under physiological conditions. rTcdA and rTcdB were then purified by Ni-NTA chromatography affinity under denaturing conditions. The rTcdA and rTcdB proteins were not found in the raw protein fraction, the flowthrough, or the eluted fractions (S2 Fig). These results suggest that we could not purify rTcdA and rTcdB using these culture conditions probably due to a lack of *tcdA* and *tcdB* expression.

To control and optimize the production yield of *C. difficile* toxins, we replaced *C. difficile tcdA* and *tcdB* gene promoters with the ATc inducible P_tet_ promoter in AD1 and AD2 respectively. This gave rise to *C. difficile* strains AD3 and AD4 containing the promoter P_tet_ in front of the *tcdA* and *tcdB* open reading frame and a sequence encoding a His-tag in the 3’ end of each gene (S1 Fig). The recombinant AD3 (P_tet_*-tcdA-*His) and AD4 (P_tet_*-tcdB-*His) strains were used to purify rTcdA and rTcdB respectively after 4h induction by ATc using Ni-NTA affinity chromatography under native conditions. SDS-PAGE gels showed an expected band of about 308 kDa (Fig 2A, lane 4) corresponding to rTcdA and an expected band of about 270 kDa (Fig 2B, lane 4) corresponding to rTcdB. Concentrations of purified toxins correspond to estimated production yields of 0.28 mg and 0.1 mg for each liter of initial culture, for rTcdA and rTcdB respectively. Our new approach allows efficient and straightforward purification of the TcdA and TcdB toxins.

**Fig 2.**
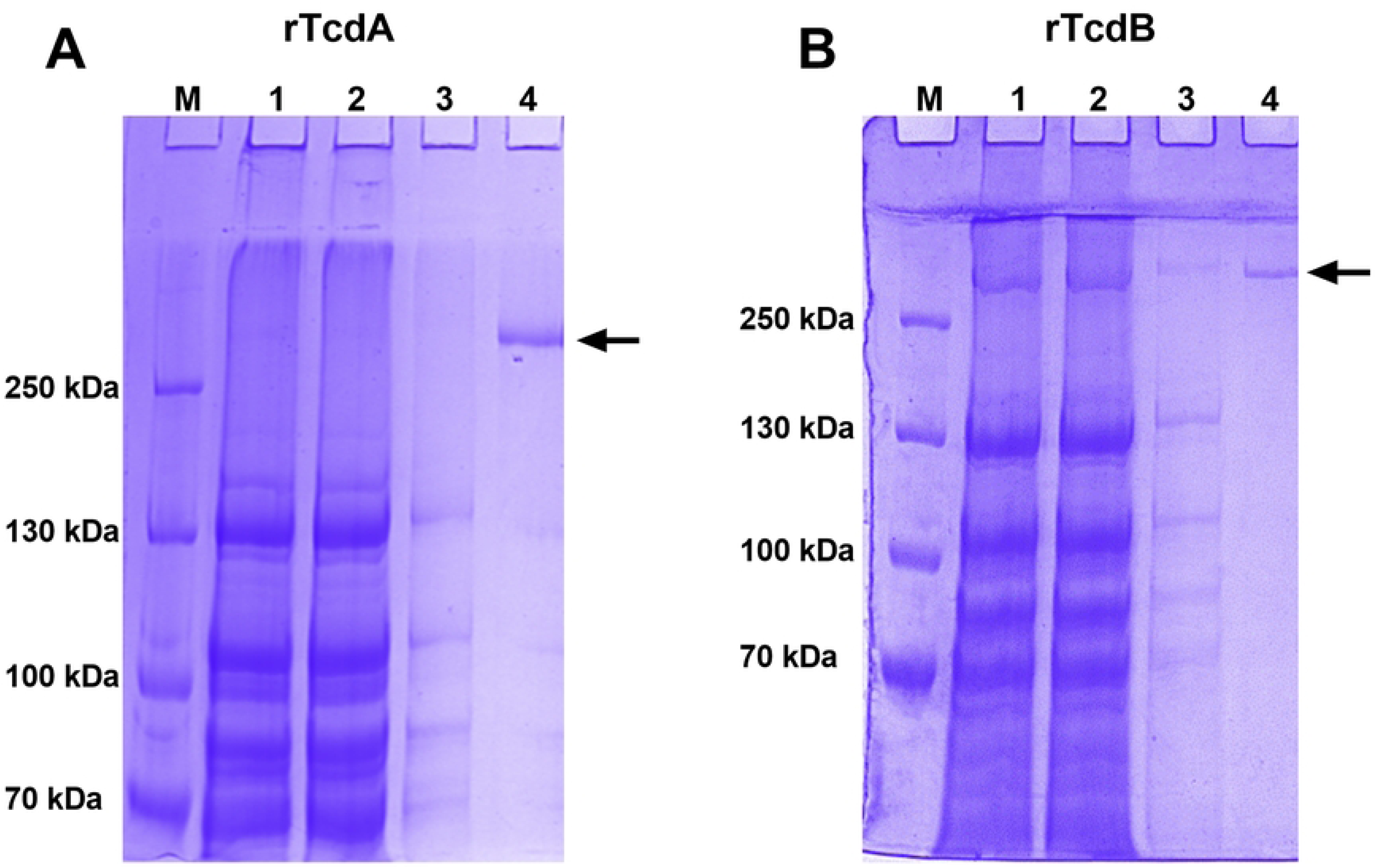
Purification of recombinant toxins by Ni-NTA affinity chromatography under native conditions. Toxins were visualized by Coomassie staining for (A) recombinant toxin A (rTcdA) and (B) recombinant toxin B (rTcdB) M: markers / 1: total bacterial lysate / 2: flow through / 3: 10 mM imidazole wash / 4: eluates after PD10. The SDS-PAGE gels shown are representative of at least 10 purifications.

### Recombinant purified toxins possess similar biological activity compared to native toxins

Functional activities of rTcdA and rTcdB were tested on Vero cells. First, we compared the cytotoxicity activity of the recombinant toxins with native toxins. After 18 hours of incubation, cells incubated with either native toxin TcdA, TcdB, purified rTcdA or purified rTcdB showed cell rounding at 4 µg.mL^-1^ (Figs 3B, 4B, 3G and 4G respectively), compared to the healthy confluent morphology observed for the negative controls (cells incubated without toxins – Figs 3A, 3F, 4A and 4F). Toxin activity decreased due to decreasing toxins concentrations, with no detectable cell rounding at 32 ng.mL^-1^ (Figs 3E, 3J, 4E, and 4J). These results suggest that rTcdA and rTcdB have similar cytotoxicity activity compared to native toxins on Vero cells and that this effect is dose-dependent.

**Fig 3.**
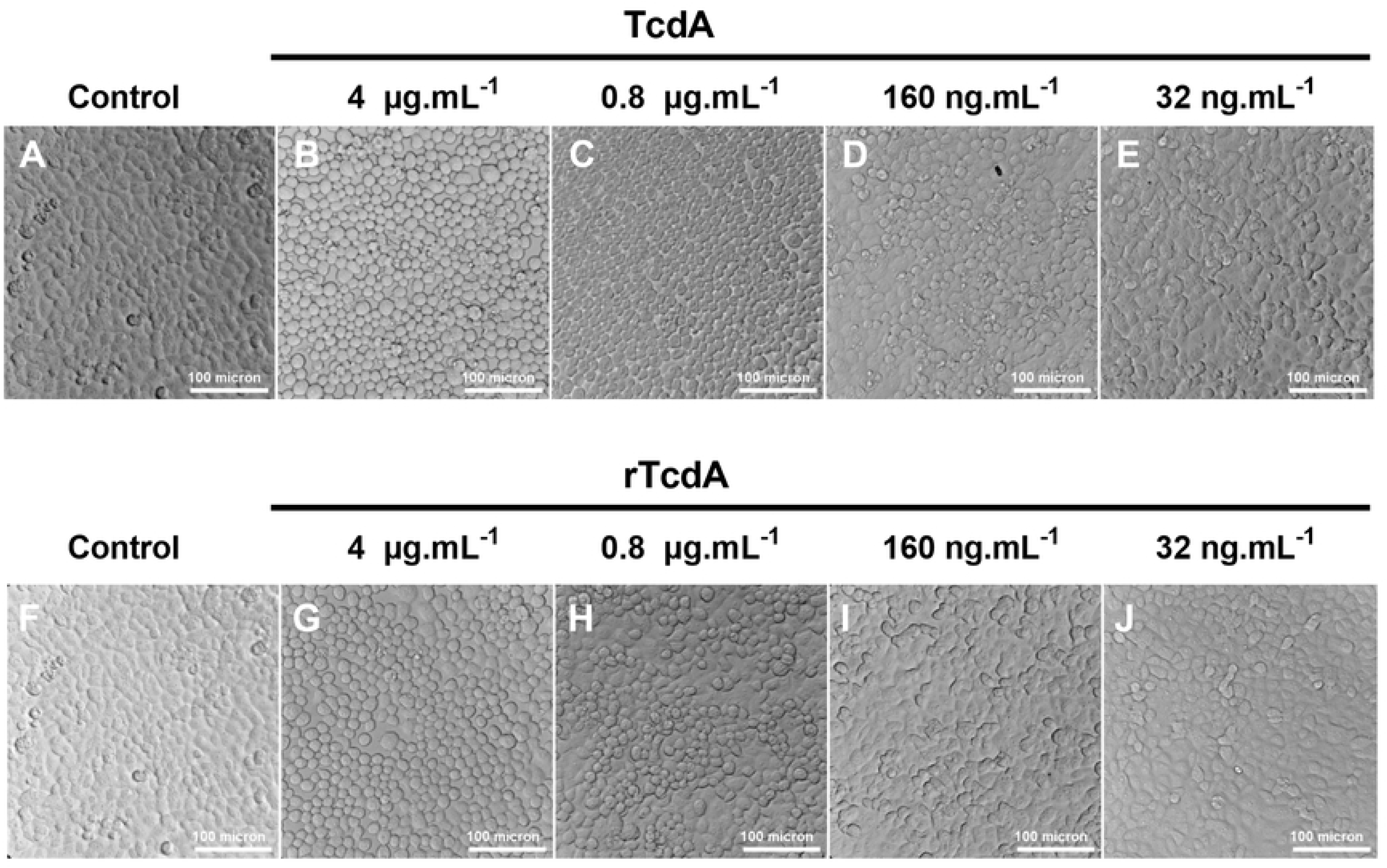
Cytotoxicity assay on Vero cells, TcdA versus rTcdA. Cells seeded in 96 wells plate were incubated in the absence of toxin (A and F) or with decreasing concentrations (4 µg.ml^-1^, 0,8 µg.ml^-1^, 160 ng.ml^-1^, and 32 ng.ml^-1^) of either native toxin TcdA (B, C, D, and E) or recombinant toxins rTcdA (G, H, I, and J). Cells were incubated for 18 hours and morphological changes were observed under phase contrast microscopy.

**Fig 4.**
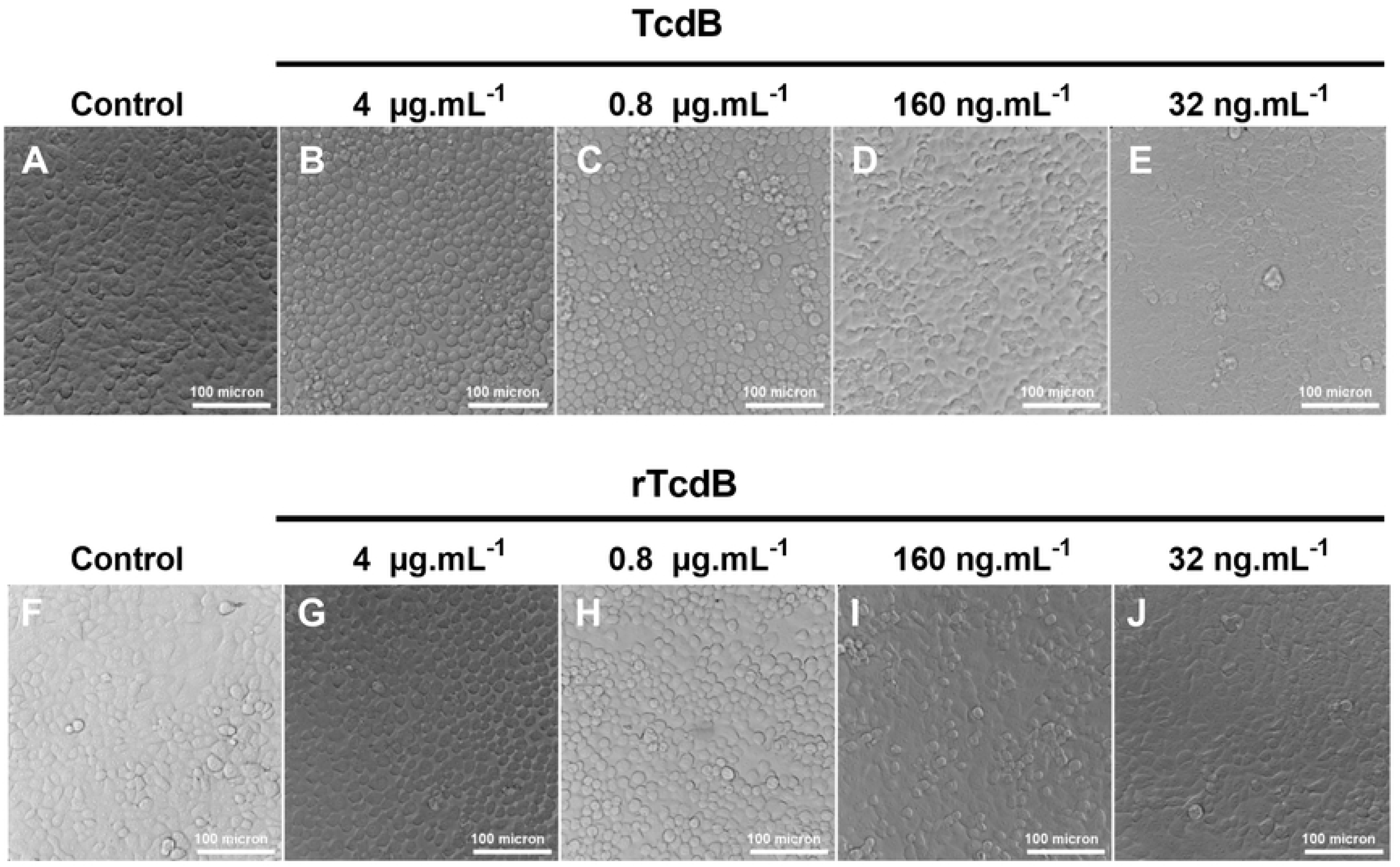
Cytotoxicity assay on Vero cells, TcdB versus rTcdB. Cells seeded in 96 wells plate were incubated in the absence of toxin (A and F) or with decreasing concentrations (4 µg.ml^-1^, 0,8 µg.ml^-1^, 160 ng.ml^-1^, and 32 ng.ml^-1^) of either native toxin TcdB (B, C, D, and E) or recombinant toxins rTcdB (G, H, I, and J). Cells were incubated for 18 hours and morphological changes were observed under phase contrast microscopy.

We then explored if purified rTcdA and rTcdB could also induce actin cytoskeleton remodeling, as described for the native toxins. Incubation of cells with either native toxins or purified recombinant toxins lead to a decrease in cell size compared to the negative control (Fig 5A and S3 Fig). Furthermore, the decrease is similar between native and purified recombinant toxins. We then looked closely at the morphological changes of the cells (Fig 5B), and we found that the actin cytoskeleton was completely disrupted with both native (Fig 5C and 5E) and recombinant toxins (Fig 5D and 5F). Taken together, these results demonstrate that the purified recombinant toxins are as active as the native toxins.

**Fig 5.**
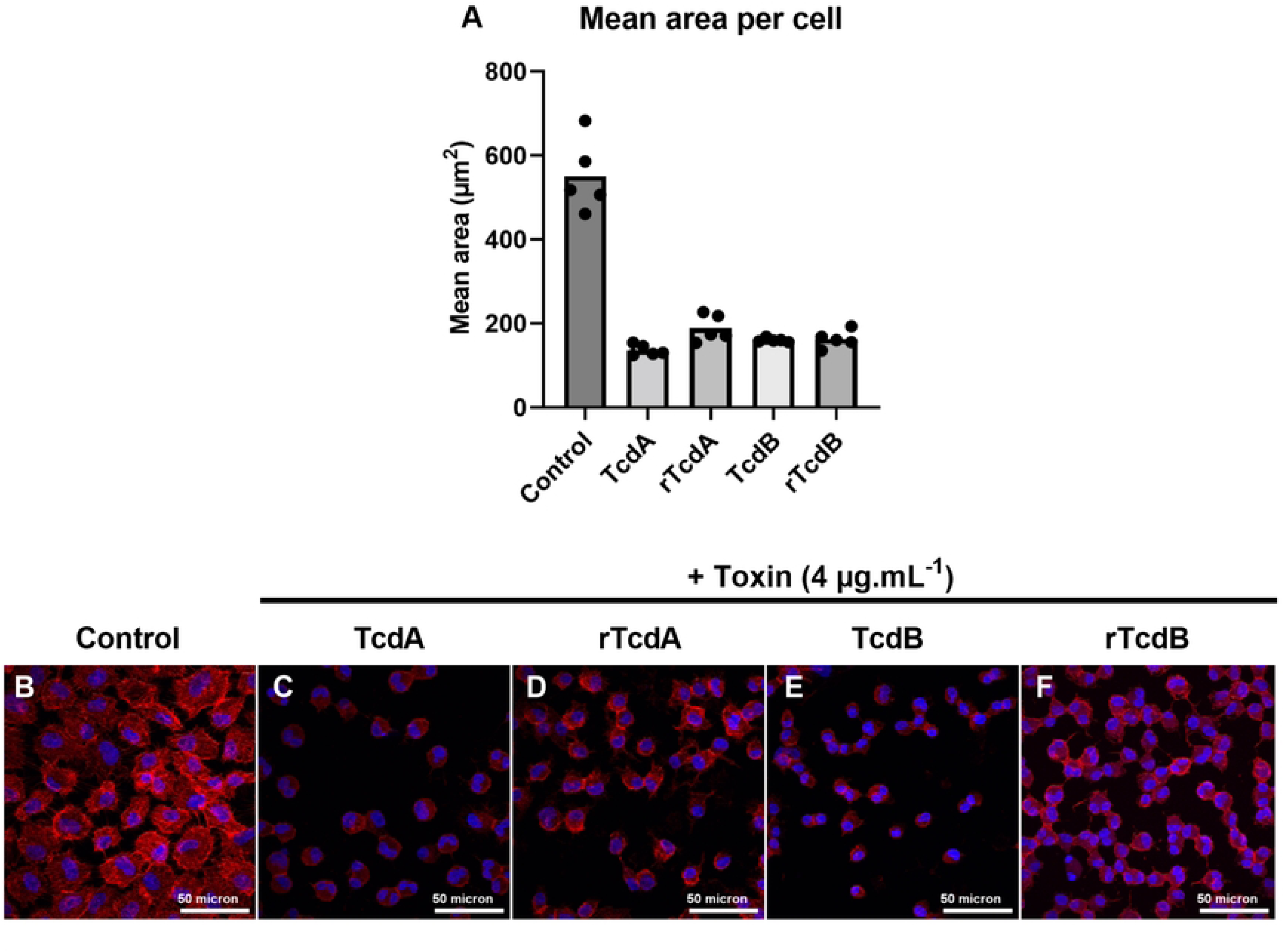
Disruption of the tight junction on Vero cells. Cells in 24 wells plate were in the absence of toxin (control) or in the presence of 4 µg.ml^-1^ of either native or recombinant toxins A and B. Cells were incubated for 18 hours, fixed, and stained with Rhodamine Phalloidin (actin staining in red) and DAPI (nuclei staining in blue) then observed under confocal microscopy with 20x objective to quantify the effect of toxins on the cells. (A) Five random fields were taken to perform an analysis with FIJI imaging software. (B-F) Cell morphology changes were observed with 63x objective, for (B) the control, (C and E) native toxins and (D and F) purified recombinant toxins at 4 µg.ml^-1^.

### Recombinant toxins can be used as substitutes for native toxins in clinical CDI investigational studies

The native TcdA and TcdB toxins are currently used in several clinical research assays, including indirect ELISA assays and neutralization antibody assays. We first wanted to evaluate the use of purified recombinant toxins as coating antigens in quantitative indirect ELISA to measure anti-TcdA and TcdB antibody titers. Thus, we compared the concentration of IgG determined for several sera from CDI recovered patients using assays done with native toxins and assays done with purified recombinant toxins. We obtained equivalent concentrations of IgG in both native and recombinant toxins assays, with a strong correlation as determined by Pearson correlation coefficient (Fig 6A and 6B).

**Fig 6.**
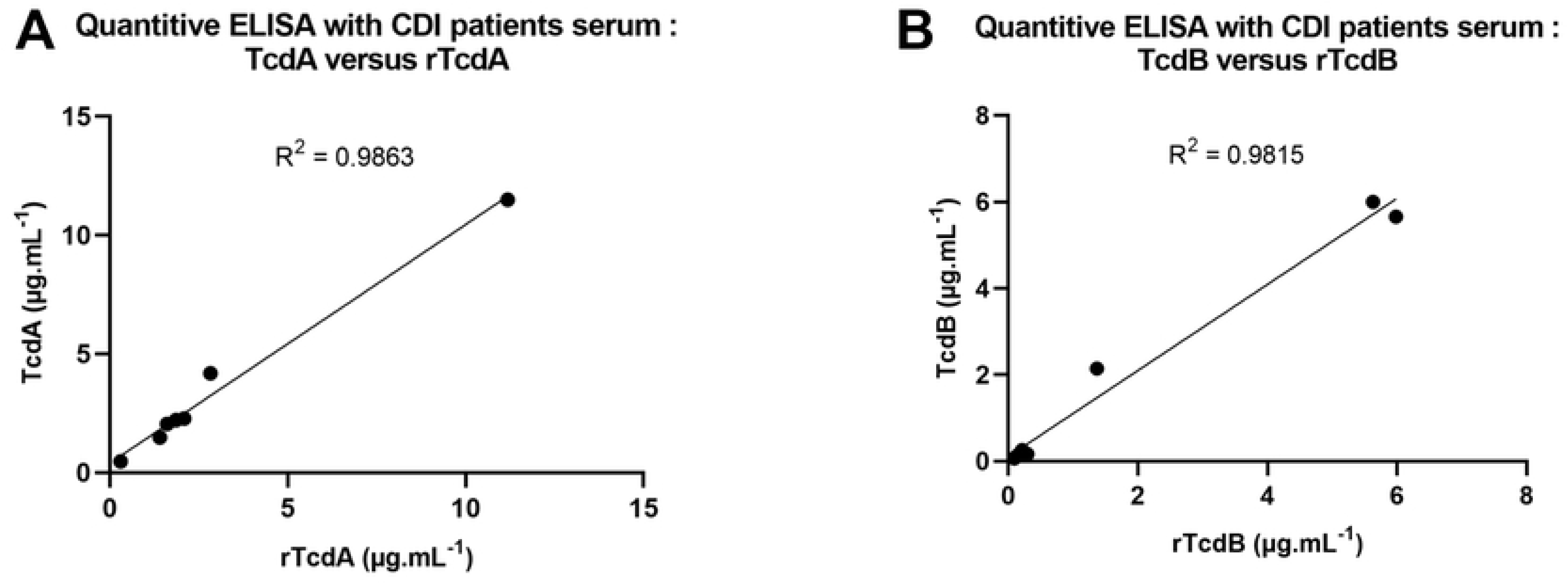
Comparison of native or recombinant toxins by quantitative ELISA. We determined the IgG concentration in 7 samples of serum from CDI patients by quantitative ELISA, for (A) TcdA or (B) TcdB. Graphs are plotted with the IgG titers obtained with recombinant or native toxins in the x and y-axis respectively. The R² represents the Pearson correlation coefficient.

Finally, we performed a neutralization antibody assay comparing both sets of toxins. In these assays, we showed that incubation of the toxins with serum samples from CDI recovered patients leads to a reduction in the cytotoxic effect of both rTcdA (Fig 7E) and rTcdB (Fig 7H). These results indicate that the effect of purified recombinant toxins can be neutralized by serum antibodies from a CDI recovered patient, confirming that the C-terminal His-tag does not hinder detection of the protein itself. Taken together, these results suggest that the purified recombinant toxins are adequate substitutes for native toxins and their use in clinical research assays.

**Fig 7.**
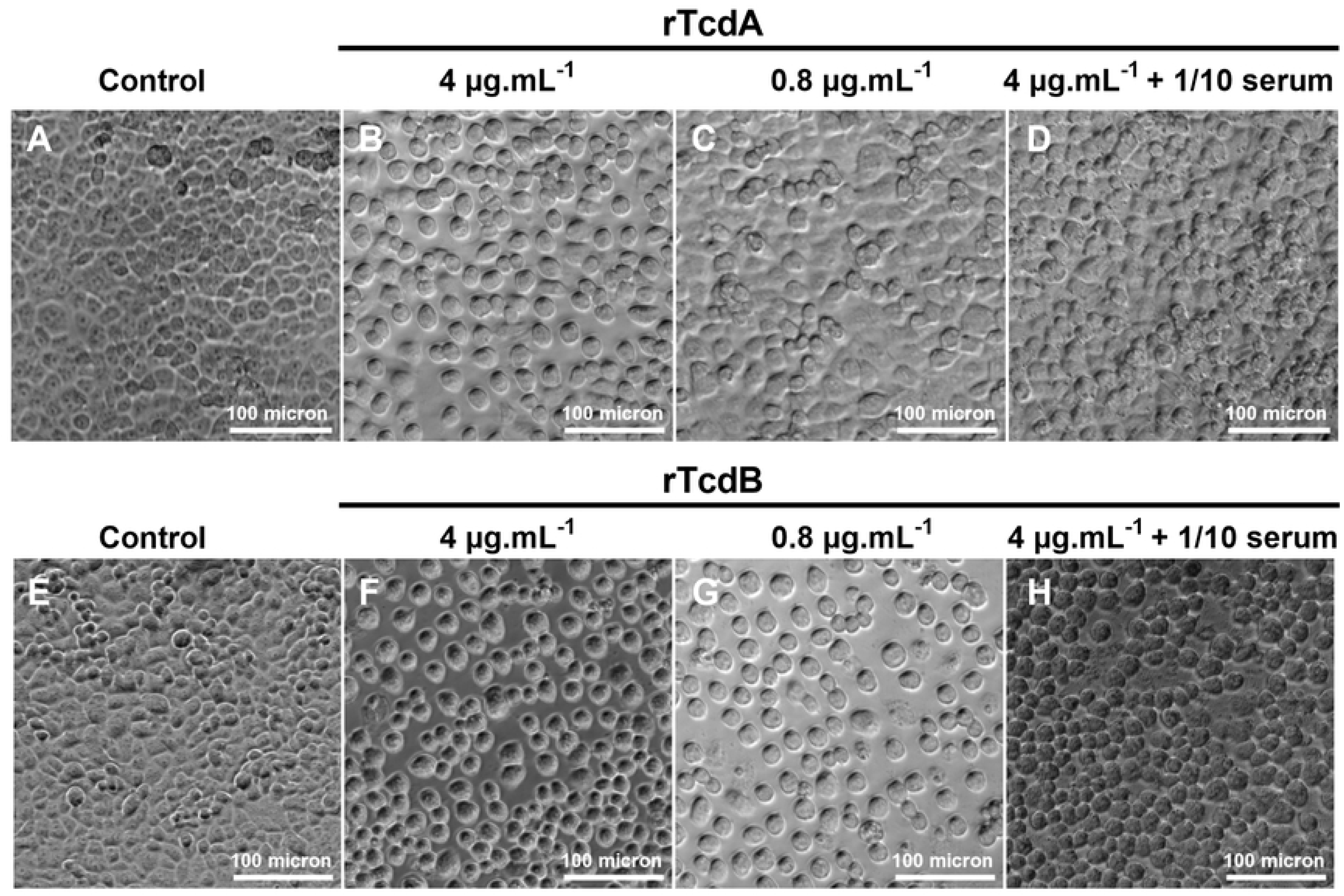
Neutralization assay on Vero cells. Cells in 96 wells plate were incubated (A and E) in the absence or with (B, C, and D) rTcdA and (F, G, and H) rTcdB at 4 µg.ml^-1^, 0.8 µg.ml^-1^ and 75 µl of 4 µg.ml^-1^ of toxins preincubated with 75 µl of 1/10 dilution of CDI patient serum. Cells were incubated for 18 hours and morphological changes were observed under phase contrast microscopy.

## Discussion and conclusion

*C. difficile* pathogenicity is linked to the production of two major virulence factors, the toxins A (TcdA) and B (TcdB). Multiple studies have increased our knowledge about the regulation, structures, and functions of *C. difficile* toxins. Recently, toxin subtypes have been identified and linked to various *C. difficile* strain clades, as well as recognition of different host receptors [16–18]. To provide new insights about their involvement in the pathophysiology of *C. difficile* infections, we need to have readily available biologically active toxins. Purified TcdA and TcdB have been useful research tools to access the host immune response against CDI by detecting the antibodies titers by ELISA assays [39–41]. TcdA and TcdB have also been used in immunization assay for the production of anti-TcdA and TcdB antibodies required in immunoblotting. Their active form have been used to study the cellular effects of these toxins in cytotoxicity assay [42], moreover in neutralization assay to study the neutralizing potential of the antibodies produced by CDI patients [31,32].

Previous purification methods of TcdA and TcdB in their native forms share similarities with those used for the purification protocols of toxins produced by other *Clostridium spp.* (*C. septicum*, *C. sordellii*, *C. perfringens*) [43–47]. In contrast with our new purification method, the purification process of native toxins is laborious and time-consuming as it requires multiple steps such as four to five days of growth in medium, ammonium sulfate precipitation and ion exchange chromatography followed by gel filtration chromatography.

To simplify this process and obtain biologically active toxins, generating recombinant toxins in a heterologous host seemed to be a viable strategy to adopt. Over the years, in the case of *C. difficile* toxins, multiple attempts have been made to purify TcdA and TcdB in *E. coli* strains, an expression system mostly used to produce recombinant proteins. Because of the large size of the *tcdA* and *tcdB* genes, expression of the entire toxin genes in *E. coli* is time-consuming and traditionally consisted in the reconstruction of cloned fragments [48]. Moreover, purification of the full-length toxin in *E. coli* was only shown for TcdA. Other studies reported the purification of fragments of TcdA or TcdB [49]. The folding of such a toxin is therefore not guaranteed. Purifying toxins from the original host is more adequate to study their structure or potential post-translational modifications.

Besides *E. coli*, Burger *et al.* and Yang *et al.* have successfully expressed and purified biological active recombinant TcdA and TcdB using *Bacillus megaterium* [50,51]. We asked for the recombinant *B. megaterium* strains expressing TcdA and TcdB. However, we could not obtain adequate titers of recombinant toxins due to difficulties in the lysis process. Finally, there are other alternatives to obtain these toxins such as commercial toxins but they are expensive and do not guarantee the biological activity. Because of these limitations and our need to use active toxins in our studies, we aimed to develop a method that simplifies the purification process of *C. difficile* TcdA and TcdB by directly using *C. difficile* strain 630Δ*erm* as the host.

The advantage of our method is, after optimizing the production and purification process, we were able to shorten toxin expression time by 10 (a half day instead of five days) and simplify the purification process compared to native toxins production. Furthermore, using the 630Δ*erm* as a host strain allows for flexibility in genetic engineering. For example, it allows the potential modification of the toxin produced, for instance to purify truncated or mutated versions of either toxin. Further work can also be done to optimize purification yields, for example by adjusting the growing media.

Using human sera from CDI recovered patients, we confirmed that these recombinant toxins could be used as antigens for ELISA for our studies based on a high correlation between the quantitative results of anti-TcdA and anti-TcdB antibodies obtained with recombinant and native toxins. In addition, our purified recombinant toxins conserved their biological activities compared to native toxins. Indeed, using cell cytotoxicity assay on Vero cells, we observed a dose-dependent cytopathic activity for both recombinant toxins and this activity can be neutralized using antibodies from a CDI recovered patient. Moreover, both recombinant toxins alter the structure of intestinal epithelia by modifying the actin cytoskeleton and opening tight junctions as already described [14,52,53].

In this study, we aimed to develop a method that simplifies the purification process of *C. difficile* TcdA and TcdB by directly using *C. difficile* strain 630Δ*erm* as the host. This allows easier genetic engineering of the strains for the optimization of the purification process, enabling other researchers to refine the methods for their own research needs. This is particularly crucial as new data emerges on the subtypes of TcdA and TcdB toxins and their respective roles. Indeed, it now appears that we can no longer consider TcdA and TcdB as two individual toxins homogenous across strains, but rather as groups of toxin subtypes with variations of structures and biological targets. For instance, TcdB2 and TcdB4 toxin subtypes have been shown to be recognized by the TFPI receptor, while TcdB1/TcdB3/TcdB5 appear to recognize Wnt receptor Frizzled proteins (FZD). Since toxins from the 630Δ*erm* or VPI 10463 strains appear in distinct clusters from toxins of the R20291 strain (Subtypes A1 / A2 and B1 / B2 respectively), toxins purified from the 630Δ*erm*/VPI 10463 strains would not be adequate proxies for the study of toxins from R20291 strain or any strain possessing TcdB2 toxins. As such, researchers interested in deciphering the roles of TcdA and TcdB will require tools that allows for the construction, production and characterization of specific toxin subtypes. To the best of our knowledge, such specificity in toxin sources and production was unattainable until now. Through this work, we successfully generated recombinant strains able to produce recombinant toxins A and B of *C. difficile* on demand, and showed that the recombinant toxins obtained have similar biological activities compared to native toxins. Simple genetic engineering is therefore all that is needed for the production of toxins in the 630Δ*erm* strain background, from the production of TcdA and/or TcdB from any *C. difficile* strain, to the construction truncated or chimeric toxins. We believe that these purified toxins and genetically engineered strains will be valuable and helpful tools used to better understand the pathogenesis of *C. difficile* infections.

## Acknowledgments

We thank J. Peltier for providing the knowledge on the obtention of the mutant by allele exchange in *Clostridioides difficile*, P. Monassa and L. Henger for the construction of plasmids pADS1 and pADS2. We also thank Pr. M. Popoff (Pasteur Institute) for generously providing *C. difficile* native toxins. We are also grateful to V. Nicolas (platform MIPSIT of Paris Saclay University) Finally, we thank our colleagues for their insightful review of the manuscript, particularly J. Malet-Villemagne who also provided us with the pJV7 plasmid. Finally, we thank C. Janoir, whose laboratory this work was conducted in, for her constant interest and support.

## Supporting information

**S1 File Protein comparisons for TcdA and TcdB from the 630*Δerm*, VPI10463 and R20291 *C. difficile* strains:** Protein identity matrixes, full-length alignment of TcdA, full-length alignment of TcdB. Protein alignments were generated with MUSCLE and visualized using BioEdit, with dots representing identical aminoacids. Protein identity matrixes were generated using BioEdit.

**Fig. S1 Strain construction for the production of the recombinant toxins rTcdA and rTcdB of *C. difficile*.** The strains were obtained in two steps using the allele exchange method. Strains ADS1 and ADS2 were obtained by adding a His-tag at the 3’ end of the *tcdA* or *tcdB* genes of the *C. difficile* strain 630Δ*erm*. Strains ADS3 and ADS4 were obtained by replacing the respective promoter of *tcdA* and *tcdB* by the promoter Ptet inducible to anhydro-tetracycline (ATc). AD3 and ADS4 were used for the purification of recombinant toxin A (rTcdA) and B (rTcdB).

**Fig. S2 Purification of recombinant His-tagged toxins by Ni-NTA affinity chromatography under denaturing conditions from the genetic construction without the Ptet promoter.** Toxins were visualized by Coomassie staining for (A) recombinant toxin A (rTcdA); (B) recombinant toxin B (rTcdB). M: markers / 1: raw protein samples / 2: flow through / 3: wash. Imidazole eluted fractions: a: 10 mM / b: 20 mM / c: 30 mM / d: 40 mM / e: 60 mM / f: 80 mM / g: 100 mM / h: 200 mM / i: 1 M

**Fig. S3 Disruption of cytoskeleton on Vero cells with 20x objective.** Cells in 24 wells plate were in the absence of toxin (control) or in the presence of 4 µg.ml-1 of either native or recombinant toxins A and B. Cells were incubated for 18 hours, fixed, and stained with Rhodamine Phalloidin (actin staining in red) and DAPI (nuclei staining in blue) then observed under confocal microscopy with 20x objective to quantify the effect of toxins on the cells.

**S1 Table List of bacterial strains used in this study.**

**S2 Table List of primers used in this study.**

**S3 Table List of Plasmids used in this study.**

**S4 Table List of Genome accession numbers used for Fig 1.**

